# Spatial structure alters the site frequency spectrum produced by hitchhiking

**DOI:** 10.1101/2022.06.08.495311

**Authors:** Jiseon Min, Misha Gupta, Michael M. Desai, Daniel B. Weissman

## Abstract

The reduction of genetic diversity due to genetic hitchhiking is widely used to find past selective sweeps from sequencing data, but very little is known about how spatial structure affects hitchhiking. We use mathematical modeling and simulations to find the unfolded site frequency spectrum (SFS) left by hitchhiking in the genomic region of a sweep in a population occupying a one-dimensional range. For such populations, sweeps spread as Fisher waves, rather than logistically. We find that this leaves a characteristic three-part SFS at loci very close to the swept locus. Very low frequencies are dominated by recent mutations that occurred after the sweep and are unaffected by hitchhiking. At moderately low frequencies, there is a transition zone primarily composed of alleles that briefly “surfed” on the wave of the sweep before falling out of the wavefront, leaving a spectrum close to that expected in well-mixed populations. However, for moderate-to-high frequencies, there is a distinctive scaling regime of the SFS produced by alleles that drifted to fixation in the wavefront and then were carried throughout the population. For loci slightly farther away from the swept locus on the genome, recombination is much more effective at restoring diversity in one-dimensional populations than it is in well-mixed ones. We find that these signatures of space can be strong even in apparently well-mixed populations with negligible spatial genetic differentiation, suggesting that spatial structure may frequently distort the signatures of hitchhiking in natural populations.

## Introduction

A selective sweep reduces neutral genetic diversity at linked loci via “genetic hitchhiking”: the genetic background on which the sweep occurred increases in frequency, while others decrease (Maynard Smith and Haigh 1974). In large populations with rapid adaptation and limited recombination, hitchhiking could be the primary factor limiting neutral genetic diversity (Gillespie 2000a,b; Weissman and Barton 2012). On the other hand, in populations where most loci are unaffected by hitchhiking, the loss of genetic diversity in specific regions of the genome is the primary method for detecting recently completed selective sweeps in natural populations (Kim and Stephan 2002; *Vitti et al.* 2013; Stephan 2019). Selection scans based on hitchhiking have found widespread sweeps in many populations and inferred their properties (Li and Stephan 2006; Sabeti *et al.* 2007; Karasov *et al.* 2010; Sattath *et al.* 2011; Vitti *et al.* 2013; Garud *et al.* 2015; Smith *et al.* 2018; Stephan 2019; Hejase *et al.* 2020; Bourgeois and Warren 2021). Two properties of particular interest are the strength of selection driving the sweep and whether the sweep is “hard” (starting from a single mutation) or “soft” (starting from multiple independent mutations) (Hermisson and Pennings 2005). Inferring these quantities requires models relating them to observable patterns of genetic diversity. Even methods that simply locate swept regions based on identifying empirical outliers need to be tested by simulations (i.e., models). It is therefore important that models for hitchhiking be at least roughly accurate descriptions of the process in natural populations.

The models underlying standard sweep-finding methods assume that the population is well-mixed (Garud *et al.* 2015; Schrider and Kern 2016; Smith *et al.* 2018; Stern *et al.* 2019; Bisschop *et al.* 2021). However, natural populations typically occupy extended spatial ranges. Often in such populations, allele frequencies are fairly constant across the range; in other words, spatial structure as measured by *F*_ST_ is weak (Hartl and Clark 1997). This reflects the fact that the mixing time is short compared to the neutral coalescence time, so well-mixed models are good approximations for many aspects of neutral evolution. However, selective sweeps are necessarily very fast compared to neutral coalescence, so for many populations, sweeping alleles will be strongly affected by spatial structure even when most neutral alleles are not (Ralph and Coop 2010). Indeed, strong patterns of spatial differentiation are one of the signatures used to detect ongoing sweeps (Sabeti *et al.* 2007; Tang *et al.* 2007; Coop *et al.* 2009). Even for methods that use these patterns to detect local adaptation rather than ongoing full sweeps (reviewed in Vitti *et al.* (2013) and Bourgeois and Warren (2021)), the premise is the same: often, dispersal is fast enough relative to drift to equalize neutral allele frequencies across space, but too slow relative to selection to maintain uniform allele frequencies at strongly selected loci.

The dynamics of sweeps in spatially structured populations can be completely different from those in well-mixed populations: instead of growing logistically, the restricted competition makes them spread as much slower traveling waves (Fisher 1937). These different dynamics can produce very different hitchhiking patterns (Slatkin and Wiehe 1998; Kim and Maruki 2011; Barton *et al.* 2013). Specifically, Barton *et al.* (2013) showed that a sweep with a given selective advantage reduces genetic diversity much less in a spatially structured population than it does in a well-mixed population of the same size. This suggests that spatial structure likely needs to be included in models of hitchhiking to get even a good first approximation.

A full description of the genetic diversity of a large sample is extremely complicated. Most of this complexity arises due to the linkage disequilibrium among alleles at different loci. Fortunately, the one-locus statistics are much simpler: when variation in allele frequencies across space is negligible, they can be completely described by the site frequency spectrum (SFS), which is the number of mutations *ξ*(*f*) found at frequency *f* in a sample of the population (Hartl 2020). Since it ignores linkage disequilibrium, *ξ*(*f*) can be measured in unphased data. For this reason, many sweep-inference methods are based on the SFS (DeGiorgio *et al.* 2016; Pavlidis *et al.* 2013; Harris *et al.* 2018; Kern and Schrider 2018). These inference methods are multi-locus in the sense that they concatenate information across genomic windows, but they are still based on one-locus statistics, i.e., they do not consider linkage disequilibrium.

In this paper, we model the hitchhiking caused by a sweep in a spatially structured population, focusing on the simplest case of a one-dimensional spatial range. We use heuristic mathematical arguments and simulations to find the effect of a sweep on the expected SFS at linked neutral loci. We start by considering a completely linked locus, and then find how hitchhiking decays as we move away from the swept locus on the genome and the recombination rate increases. We find that even in weakly spatially structured populations, space creates a distinctive tail in the expected site frequency spectrum (SFS) and changes the width of the region of the genome affected by hitchhiking. We discuss the potential implications of our results for the accuracy of inferences about adaptation based on well-mixed models.

## Model

We use a one-dimensional stepping-stone model, in which there is a line of *L* demes, each with a fixed population size of *ρ* hap-loid individuals, for a total population size of *N* = *Lρ*. Within each deme, we use a Wright-Fisher model, with individuals having a probability *m* each generation of being a migrant from a neighboring deme. The migrants have an equal probability of coming from the neighboring deme to the right or the left. At the edges of the range, the leftmost and rightmost demes have only one neighboring deme each, so individuals in these demes have a probability of only *m*/2 per generation of being a migrant.

At some point in time, an individual acquires a beneficial mutation with selection coefficient *s*. Most such mutations rapidly go extinct due to drift; we discard these simulations and instead keep only those in which the mutation is lucky enough to escape extinction and sweep to fixation. For simplicity, we assume the beneficial mutation occurs in the leftmost deme in most of our results. As we discuss in Appendix C, this captures all the main qualitative patterns, and generalizing the quantitative patterns to mutations starting in other locations is straightforward. We focus on the case *s* < *m* and 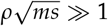 in which the traveling wave of the successful beneficial allele has a front that is aproximately continuous in space, with a characteristic width 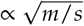 and speed 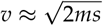 (Barton *et al.* (2013); illustrated by the black curve in Figure 1). In this approximately continuous case, we do not expect any of our results to depend on the details of the demic structure. At finite densities *ρ*, the wave speed is slightly smaller than 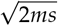 (Barton *et al.* 2013), so in all plots we use the speed measured directly from the simulations (see Appendix B for details).

**Figure 1.**
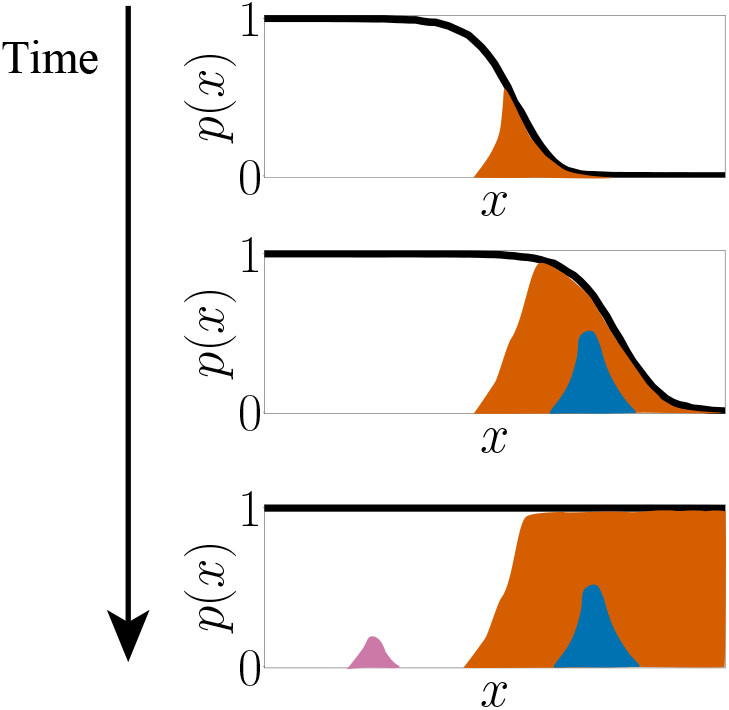
Spatial hitchhiking leaves three classes of alleles: those that fix in wavefront (orange), those that surf but do not fix (blue), and those that arise behind the wavefront (pink). The black curve shows the frequency of the beneficial allele at location *x* and each colored area represents frequency of different neutral mutant lineage.

For the hard, complete sweeps in our model, all neutral diversity after the end of the sweep must be produced by mutation or recombination. We follow a single neutral locus with mutation rate *U*_n_ and recombination rate *r* with the selected locus. We use the infinite allele model in which every new neutral mutation is unique. This is a good first approximation as long as the population size *N* times the per-base neutral mutation rate is small compared to one. Our neutral locus could extend over multiple bases, as long as recombination among them is negligible; see the Discussion for consideration of possible locus sizes.

We focus on the parameter regime in which spatial structure has a negligible effect on neutral allele dynamics in the absence of the sweep, but a strong effect on the sweep dynamics. To satisfy the first condition, we set the parameters so that the mixing time from dispersal, *T*_mix_ ∝ *L*^2^/*m*, is much smaller than the well-mixed coalescent time scale, *T*_coal_ = *N*. This makes it so that neutral heterozygosity is nearly unaffected by space, i.e, so that the “effective population size” is *N_e_* ≡ *T*_coal_ ≈ *N* (Maruyama 1971). To satisfy the second condition, we make sure that the mixing time *T*_mix_is long compared to the duration of the sweep, 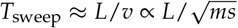. This is roughly equivalent to requiring that the range size is long compared to the wave-front of the sweep, 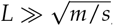, or that the mixing time be long compared to the time the sweep would take in a well-mixed population, *T*_mix_ ≫ ln(*Ns*)/*s*. We can summarize these conditions as *T*_sweep_ ≪ *T*_mix_ ≪ *T*_coal_.

In this parameter regime, common alleles at sites unlinked to the sweep have little variation in their frequencies across space. During and immediately after the sweep, variation at loci linked to the sweep may vary strongly in space (Booker *et al.* 2020), and the increased mixing caused by hitchhiking (*Barton et al.* 2013) can lead to distant individuals sharing unusually recent common ancestry near the swept locus (Allman and Weissman 2018). But dispersal will erase these spatial patterns in a time *T*_mix_ much shorter than the time *T*_coal_ that it takes drift to erase the effect of hitchhiking on the overall allele frequencies (Ralph and Coop 2010). Thus, for most potentially detectable sweeps, there will be no direct signature of the effect of spatial structure. In other words, we do not expect to see allele frequencies varying much in space, but the strong spatial structure that existed during the sweep may still leave a large signature in the overall SFS. Throughout the paper, we will focus on the unfolded SFS, assuming that there is an outgroup such that ancestral and mutant alleles can be distinguished, as the folded SFS can be immediately derived from the unfolded one.

We follow Barton *et al.* (2013) in conducting our simulations in two steps. First, we simulate the dynamics at just the selected locus forward in time, saving the trajectory of the sweep in time and space. We then use this saved trajectory to conduct structured coalescent simulations of the linked neutral locus backward in time. This increases computational efficiency because most of the stochasticity in the SFS is produced in the coalescent process rather than the sweep trajectory, and we can simulate many independent coalescent histories on each simulated sweep trajectory. We find the expected SFS by averaging our results over many independent forward simulations and many more independent backward simulations for each forward simulation. Because we focus on the expected SFS, we do not need to simulate neutral mutations explicitly but can instead simply multiply the branch lengths of our simulated coalescent trees by *U*_n_ to find the expected number of mutations. The neutral mutation rate *U*_n_ is, therefore, simply an overall scaling factor in all our results. For the details of the simulations, see Appendix A. All code can be found at https://github.com/weissmanlab/SFS_spatial_sweep_1d_Genetics.

## Results

### Background: Hitchhiking at a completely linked locus in a well-mixed population

We first review the expected SFS left by hitchhiking in a well-mixed population. Here, we focus on the simplest case of a completely linked locus, sampled immediately after the fixation of the beneficial allele; we consider the effects of drift after the sweep and recombination below. All diversity is, therefore, from mutations that occurred after the beginning of the sweep in individuals carrying the beneficial allele. The expected SFS follows two power-laws (Figure 2, grey curve). Almost all individuals carrying the sweeping allele who could potentially be ancestral to the sample lived in the late stages of the sweep, so most mutations are relatively recent and occurred too late to hitchhike to high frequencies. Their dynamics are, therefore, simply controlled by drift and follow the neutral expectation (Wakeley 2008):

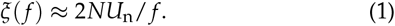

Eq. (1) is shown as a pink dashed curve in Figure 2 and following figures. Since these mutations occurred less than ln(*Ns*)/*s* generations before sampling (i.e., during the sweep), they do not have time to drift to a frequency of more than about *f* ≲ ln(*Ns*)/(*Ns*) (Desai and Fisher 2007) (Figure 2, vertical dotted turquoise line). Only the lucky older mutations that occurred when the sweeping allele was rare can hitchhike to frequencies *f* ≫ ln(*Ns*)/(*Ns*). Since the beneficial background was growing approximately exponentially at this time, the frequency spectrum of mutations follows the classic scaling ∝ 1/*f*^2^ of Luria and Delbrück (1943). Specifically, for a mutation to hitchhike to frequency of at least *f*, it must have occurred when the number of individuals with the beneficial allele was ≲ 1/*f*. The total number of mutational opportunities up to this point of the sweep is equal to the number of individuals with the beneficial allele integrated over time, ≈1/(*sf*), so the expected number of mutations with frequency > *f* is ≲ *U_n_*/(*s f*). Differentiating the expected number of mutations with respect to frequency gives the SFS:

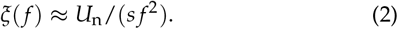

Eq. (2) is shown as a dot-dashed cyan curve in Figure 2.

**Figure 2.**
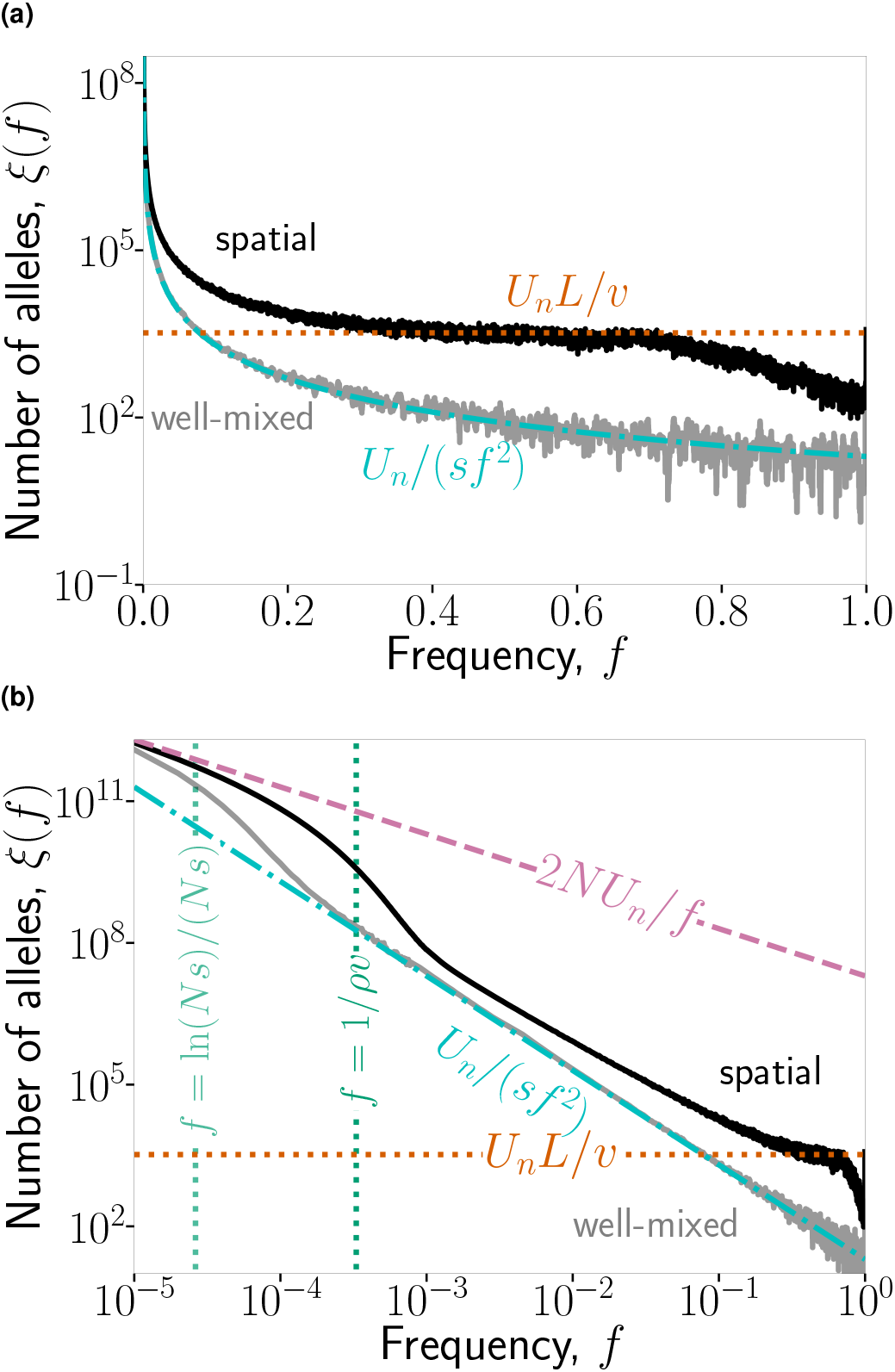
Spatial structure creates a distinctive high-frequency tail in the site frequency spectrum (SFS) left by hitchhiking. Black and grey curves show simulation results for the SFS at a neutral locus completely linked to a selective sweep in a one-dimensional and well-mixed population, respectively, with the same population size (*N* = 10^7^) and selective coefficient (*s* = 0.05). **(a)**: For high-frequency polymorphisms, the spatial SFS is flat, following *ξ*(*f*) ≈ *U*_n_ *L*/*v* (dotted orange line), and much higher than the well-mixed SFS, *ξ*(*f*) ≈ *U*_n_/(*s f*^2^) (dot-dashed cyan curve). **(b)**: Plotting frequency on a log scale shows that at very low frequencies, both spectra match the neutral expectation *ξ*(*f*) = 2*NU*_n_/*f* (pink dashed line). For frequencies *f* ≫ ln(*Ns*)/(*Ns*) (turquoise vertical line), the well-mixed SFS follows the high-frequency law. The spatial SFS deviates from neutrality at a higher frequency, *f* ≫ 1/(*ρv*), and follows a slightly different law—see Figure 3. The populations were sampled immediately after the fixation of the beneficial allele. Other model parameters: *U*_n_ = 1, *m* = 0.25, *ρ* = 2 × 10^4^, *L* = 500. See Appendix A for details of how simulation results are averaged over multiple runs and frequencies in this and following figures.

### Completely linked locus in a one-dimensional population

We will start by sketching an intuitive argument for the qualitative features of the SFS left by hitchhiking in a one-dimensional population. As above, we will first focus on a completely linked locus sampled immediately after the sweep. We do not attempt to characterize spatial patterns in genetic diversity at the locus even though we consider just-completed sweeps because we are interested in more typical older sweeps, finished at least ≈*T*_mix_ generations ago. In a one-dimensional population, the sweep proceeds as a Fisher wave (Fisher 1937) (Figure 1, black curves). Most neutral mutations ancestral to the sample will occur in the bulk of the wave, where the beneficial allele is already fixed and there is no more hitchhiking (Figure 1, pink). These mutations should follow the neutral SFS and should be limited to low frequencies (Figure 2, pink dashed line). Yet, the mutations that occur within the wavefront can hitchhike and follow either of two possible trajectories. First, most will “surf” in the wavefront temporarily, but eventually be left behind and stop hitchhiking (Figure 2, blue). The luckiest mutations will manage to fix in the wavefront and hitchhike to very high frequencies as they are carried by the wave of the beneficial allele (Figure 2, orange). We, therefore, expect to see three basic regimes in the hitchhiking SFS of a one-dimensional population, rather than just the two regimes in the well-mixed population. This intuition is confirmed by simulations (black solid curve in Figure 2). These show that the expected SFS of a one-dimensional population is higher than the one of a well-mixed population, i.e., there is more genetic diversity.

We now give approximate analytical expressions for the form of the SFS in the three different regimes. Since the low-frequency alleles that occurred in the bulk of wave (Figure 1, pink) did not hitchhike, they simply follow the neutral expectation *ξ*(*f*) ≈ 2*NU*n/*f* (Figure 2, pink), as in a well-mixed population. But since the sweep time is now much longer than it would be in a well-mixed population, *T*_sweep_ ≈ *L*/*v* ≫ ln(*Ns*)/*s*, these mutations can drift for longer and reach higher frequencies, *f* ≲ *T*sweep/*N* ≈ 1/(*ρv*) (Figure 2, vertical green dotted line).

To reach higher frequencies, *f* ≫ *T*_sweep_/*N*, mutations have to hitchhike. For most of this frequency range, the SFS is approximately uniformly distributed, a pattern unique to the spatially extended population. This tail reflects the mutations that succeed in fixing in the wavefront (Figure 2, orange). Since the dynamics within the front are neutral, mutations fix in the front at a steady rate of *U*_n_. For such a mutation to reach frequency at least *f*, it must have fixed in the wavefront before the sweeping allele reached frequency 1 − *f*. The total amount of time that sweeping allele spends at frequency less than 1 - *f* is ≈ (1 - *f*)*L*/*v*, so the total number of mutations that we expect to fix in the front and reach frequencies greater than *f* is *U*_n_(1 - *f*)*L*/*v*. Differentiating with respect to *f* gives the expected SFS:

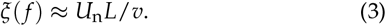

This analytical approximation is in good agreement with simu-13 lations (Figure 2, orange horizontal dotted line). It breaks down 14 at very high frequencies (*f* ≈ 0.8 in Figure 2), at which point the 15 initial waiting time for the beneficial allele to settle down to a 16 Fisher wave becomes important; see Appendix D.

Between the low-frequency neutral regime and the high-frequency flat regime described above, there can also be an intermediate-frequency hitchhiking regime (frequencies ~10^−3^ to ~10^−1^ in Figure 2). This regime consists of mutations that occurred in the wavefront in the sweeping background and surfed for a while but were then left behind (Figure 1, blue area). The expected SFS of a one-dimensional population in this frequency range is similar to but distinct from that of a well-mixed population with the same *N* and *s* (*ξ*(*f*) ≈ *U*_n_/(*s f*^2^), as described above). Intuitively, the similarity is because these mutations only surf over a small portion of the range, so they have little opportunity to “feel” the spatial structure. To characterize the distinctive behavior of the one-dimensional spectrum in this regime, we can treat the wavefront as its own small population with a coalescent time scale *T*_front_ given by the power-law fits of Barton *et al.* (2013) and Birzu *et al.* (2018). Coarse-graining over this time scale then produces an effective offspring distribution with a heavy-tailed “sweepstakes” pattern (Okada and Hallatschek 2021). This gives a site frequency spectrum:

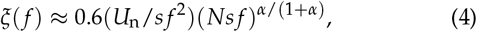

with *α* ≈ 0.3 (Barton *et al.* 2013); see Appendix E for details. The prefactor of 0.6 is a fit to the simulations shown in Figure 3. Because mutations can only persist in the wavefront for a limited amount of time before falling behind or fixing, this regime can only cover a limited frequency range that shrinks in longer habitats. This regime is also the first to be eroded as time passes from the end of the sweep (Figure 4). Taken together, these factors suggest that the higher-frequency uniform regime may be the more relevant portion of the spectrum for data from natural populations.

### The signature of spatial structure persists through time

Above, we have assumed that the population is sampled immediately after the sweep, which is unusual in reality. More often, one is interested in all the sweeps that may have happened recently enough to still be detected. Since the signature of the sweep lasts much longer than the sweep itself, the waiting time *T*past between the end of the sweep and sampling is typically longer as well: *T*_past_ ≫ *T*_sweep_. Intuitively, this waiting time allows more time for new mutations to appear and grow neutrally via genetic drift; therefore, it extends the part of the SFS that matches the neutral expectation *ξ*(*f*) ≈ 2*NU*_n_/*f*. More specifically, since drift has ≲ *T*_sweep_ + *T*_past_ generations to act, the neutral part of the spectrum extends to *f* ≲ (*T*_sweep_ + *T*_past_)/*N*, until drift erases the entire signature of hitchhiking at *T*_past_ ≈ *N*. For mutations at much higher frequencies *f* ≫ (*T*_sweep_ + *T*_past_)/*N*, the change in frequency due to drift is minor, so this part of the spectrum is nearly unaffected. (The exception to this is the extreme high-frequency tail where the frequency of the ancestral allele is small, 1 - *f* ≲ (*T*_sweep_ + *T*_past_)/*N*; these mutations can be driven to fixation by drift.) Figure 4 shows that this intuition matches simulations. As a result, the high-frequency uniform tail of the hitchhiking SFS in one-dimensional populations is the longest-lasting part of the spectrum, while the intermediate-frequency regime relaxes to neutrality first.

**Figure 3.**
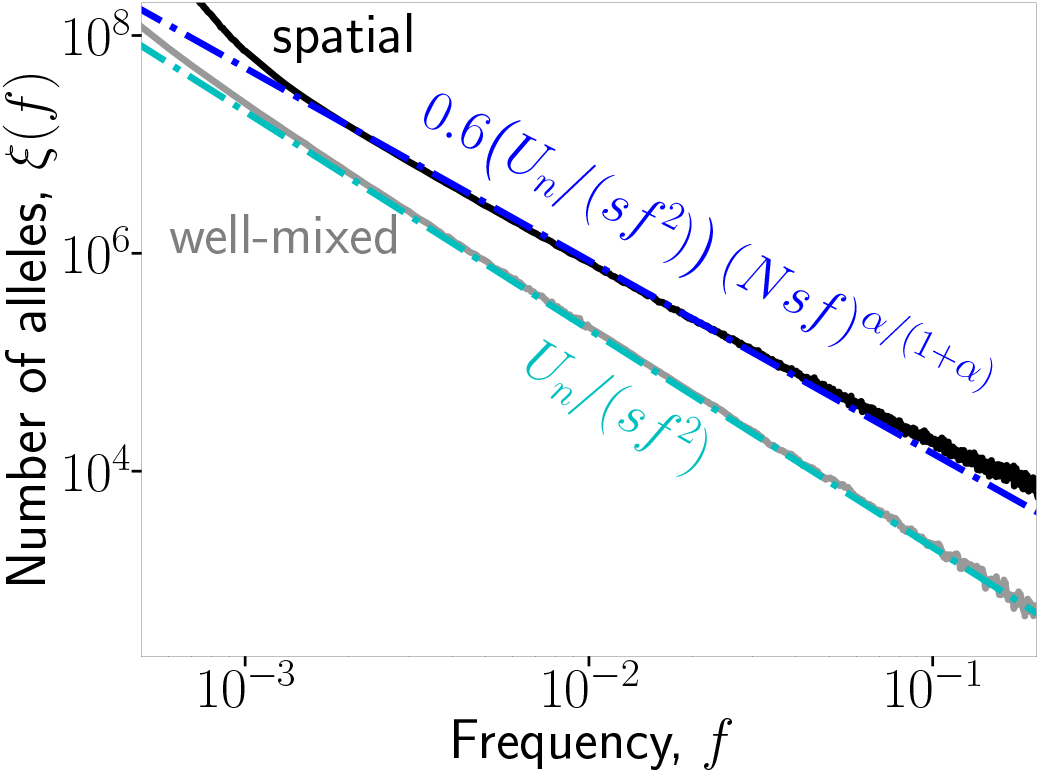
In spatial populations, alleles that temporarily surf in the wavefront create an intermediate-frequency regime in the SFS. Plot shows a close-up on the intermediate frequencies for the same data shown in Figure 2. The blue dot-dashed curve shows that the spatial SFS in this regime is different from that of the well-mixed one by a factor ≈ 0.6(*Ns f*)^*α*/(1+*α*)^, with *α* ≈ 0.3.

**Figure 4.**
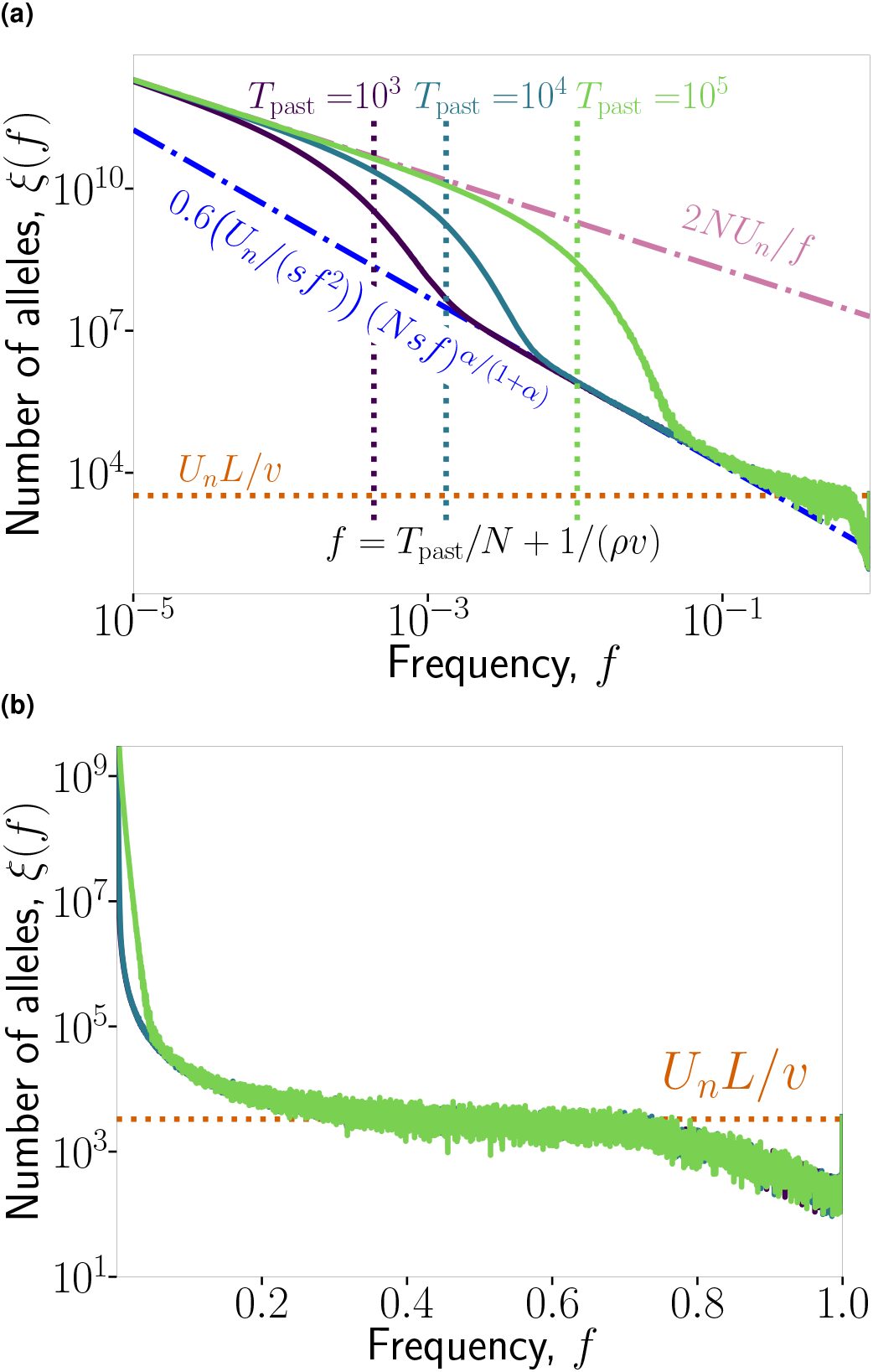
The distinctive high-frequency tail of the SFS persists long after the sweep is completed. The purple, teal, and green curves show simulation results for the expected SFS left by hitchhiking *T*_past_ = 10^3^, 10^4^, and 10^5^ generations after the beneficial allele fixes, respectively. The vertical dotted lines of the corresponding colors show the predicted upper extent *f* = (*T*_past_ + *T*_sweep_)/*N* of the approximately neutral part of the SFS for each value of *T*_past_. Parameters: *L* = 500, *ρ* = 2 × 104, *m* = 0.25, *s* = 0.05, *U*_n_ = 1, *α* = 0.30.

### The effect of recombination

So far, we have focused on neutral loci that are completely linked to the positively selected locus. We now consider a neutral locus farther away on the genome that recombines at rate *r* with the selected locus. We can do this by combining the picture of the one-dimensional dynamics developed above with the known effects of recombination on the SFS produced by hitchhiking. In this section, we will first review those effects and their mathematical expressions in well-mixed populations. Then we will introduce the equivalent expressions for one-dimensional populations. Recombination has two related effects: first, it brings new alleles onto the sweeping background, allowing them to hitchhike, effectively increasing the mutation rate; and second, it breaks down the positive linkage disequilibrium that drives the hitchhiking. While these effects are really two sides of the same coin, it is helpful to think of them separately because they vary in importance at different genomic scales.

For the first effect of recombination, bringing new alleles onto the sweeping background, a typical recombinant will be very distantly related to the original sweeping background, with an expected time to the most recent common ancestor of *T*_coal_ ≈ *N*. They will therefore differ by an average of 2*NU*_n_ mutations, *NU*_n_ each on the original background and the recombinant. The fact that the recombinant brings in *NU*_n_ new mutations means that the effective neutral mutation rate of the sweep is increased by *rNU*_n_, for a total of *U*_n_,eff = (1 + *Nr*)*U*_n_. On the other hand, the fact that the recombinant has the ancestral allele at *NU*_n_ sites where the original sweeping background was mutated, means that those mutations are prevented from fixing by the hitchhiking of the *ancestral* alleles. (In the absence of recombination, the mutant alleles would have fixed and contributed to divergence from an outgroup but not polymorphism.) This produces a mirror uptick in the unfolded SFS at high frequencies, where at effective mutation rate *NrU*_n_, ancestral alleles arrive on the sweeping background and hitchhike following the same expressions as previously found for the mutant alleles, but with the ancestral allele frequency 1 - *f* replacing the mutant frequency *f*. (From a coalescent perspective, the two terms are both produced by the same dynamics—a set of lineages coalesces during the sweep but their ancestor recombines out before the sweep origin—with the only difference being which branch of the tree a mutation falls on.) Thus, in a well-mixed population, Eq. (2) for the SFS becomes:

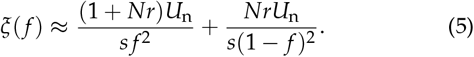

As with a completely linked locus, Eq. (5) is only valid for allele frequencies *f* above that which can be reached by drift. It is also cut off at an upper limit of *f* ≈ 1 - (*r*/*s*) ln(*Ns*), above which the SFS flattens out (Fay and Wu 2000). Within these bounds, Eq. (5) matches simulations well, although it slightly underestimates the rate at which the SFS relaxes to neutral expectation for high recombination rates (Figure 5a-c, cyan dot-dashed lines). We see that this first effect of recombination is important even very close to the selected locus, with recombination dominating the effective mutation rate *U*_n_,eff for *r* ≳ 1/*N*, and the high-frequency mirror-image portion of the spectrum being visible at even closer loci in large samples.

**Figure 5.**
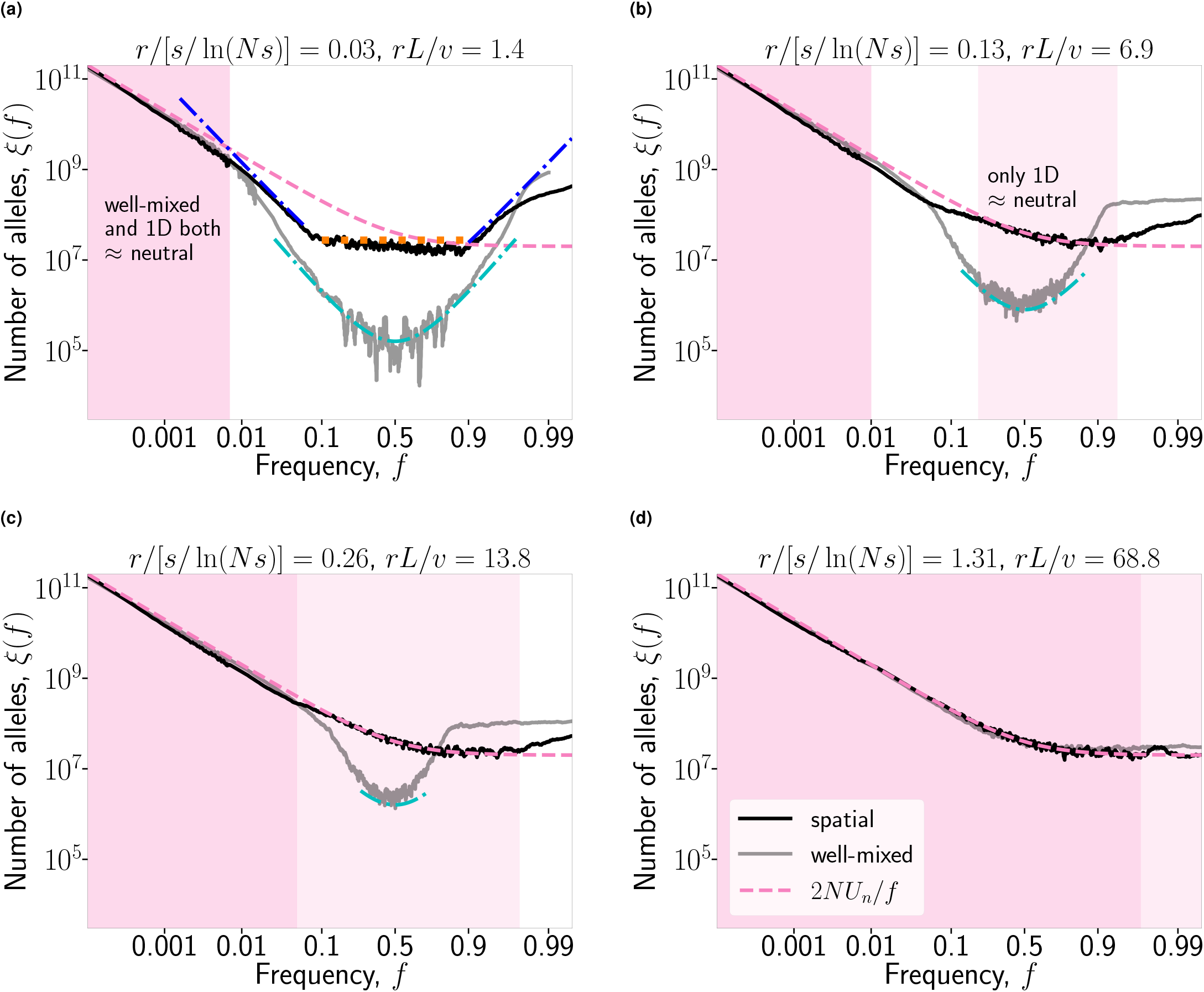
Recombination limits hitchhiking more in a one-dimensional population than in a well-mixed one. Panels show the effect of varying recombination rate *r* on the SFS while keeping other parameters fixed. In each panel, the population size *N* = 10^7^, the selection coefficient *s* = 0.05, and the recombination rates *r* are the same for the simulated one-dimensional (black) and well-mixed (grey) populations. The dark pink shaded regions illustrate the frequency ranges where both populations’ spectra are close to the neutral expectation, while the light pink illustrate where only the one-dimensional population is. When *r* falls in between the scales set by the sweep time in a one-dimensional population and the sweep time in a well-mixed population, *v*/*L* ≪ *r* ≪ *s*/ln(*Ns*), hitchhiking is weak in the one-dimensional population but strong in the well-mixed one. For the one-dimensional population, *L* = 2000, *ρ* = 5000, and *m* = 0.25. The pink dashed line shows the neutral SFS (*ξ*(*f*) = 2*NU*_n_/*f*) as in previous figures. The orange dotted line and blue and cyan dot-dashed lines are similar to those in previous figures but adjusted for the effects of recombination; see Eq. (6), Eq. (7), and Eq. (5), respectively. Populations were sampled immediately after the end of the sweep.

The second effect of recombination, reducing hitchhiking by breaking down positive linkage disequilibrium between the sweeping allele and alleles at neutral loci, is more familiar than the effect described above, but it is only effective at more distant loci. Intuitively, this reflects a comparison between the time scale for hitchhiking to increase the frequency of a neutral allele in positive linkage disequilibrium with the sweeping allele, and the time scale 1/*r* for the linkage disequilibrium to decay. For a well-mixed population, the time for hitchhiking to boost an allele initially established at ≈1/*s* copies to frequency *f* is ≈ln(*N f s*)/*s*. Thus, recombination prevents hitchhiking from affecting the bulk of the SFS at loci map distance *r* ≳ *s*/ ln(*Ns*) from the sweep (Stephan *et al.* 1992; Kaplan *et al.* 1989; Barton 1998, 2000). The effect on the upper tail of the SFS persists out to slightly more distant loci because it involves alleles that hitch-hike for a slightly shorter time and because the neutral spectrum is lowest there (Fay and Wu 2000); this is barely visible in Figure 5d, but even this is effectively gone before the recombination rate reaches *r* ≈ *s*.

In a one-dimensional population, the logic is exactly the same, except that two expressions, Eq. (3) and Eq. (4), need to be adjusted. For the alleles that fix in the wave front, Eq. (3) becomes:

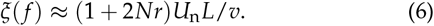

The factor (1 + 2*Nr*)*U*_n_ is the sum of mutations arising during the sweep (*U*_n_), mutations that recombine onto the sweeping background (*NrU*_n_), and ancestral alleles that replace mutations when they recombine onto the sweeping background (another factor of *NrU*_n_). Since there is no *f* dependence, these all combine into one prefactor, rather than appearing in front of separate terms as in Eq. (5). Eq. (6) matches simulations well (Figure 5a, orange dotted line). The factor *rL*/*v* in Eq. (6) is the expected number of recombinant lineages that fix in the front during the sweep, and thus also appears (inversely) in the expression (C.3) in Barton *et al.* (2013) for the probability of coalescence of a pair of lineages due to hitchhiking.

For the alleles that only surf temporarily in the front before being dropped, Eq. (4) becomes:

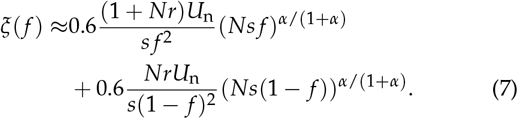

As in Eq. (5), the first term comes from the hitchhiking of mutations, while the second comes from the hitchhiking of ancestral alleles preventing the fixation of hitchhiking mutations. Eq. (7) matches simulations well, but only applies over small ranges of frequencies (Figure 5a, blue dot-dashed lines); as in the well-mixed case, it does not apply to very low or high frequencies where drift becomes more important, and it also does not apply to the middle of the frequency range, where Eq. (6) dominates.

In a one-dimensional population, sweeps are slower than in a well-mixed population. Therefore, smaller recombination rates *r* are sufficient to produce linkage disequilibrium decay times 1/*r* that are short compared to the time needed to hitchhike, the necessary condition for the second effect of recombination (i.e., recombination blocks hitchhiking). The one-dimensional population differs from the well-mixed one in that there are now two different hitchhiking time scales, corresponding to the two regimes of the completely linked SFS: the long time *T*_sweep_ ≈ *L*/*v* over which alleles that fix in the front hitchhike, and the shorter time 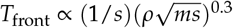 (Barton *et al.* 2013; Birzu *et al.* 2018) over which alleles that only temporarily surf in the front can hitchhike. As we move away from the swept locus on the genome, we expect the uniform portion of the hitchhiking SFS Eq. (6) to relax to neutrality first, at *r* ≈ *v*/*L*, while the intermediate-frequency signal persists until map distances 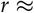 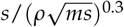 that are similar to those at which intermediate frequency signal vanishes in well-mixed populations.

## Discussion

We have shown that the hitchhiking caused by a selective sweep in a one-dimensional spatially structured population produces a very different expected site frequency spectrum from that left by hitchhiking in well-mixed populations. This is true even if the spatial structure is very weak, in the sense that frequencies of common neutral alleles vary very little from location to location. The most striking feature of the expected one-dimensional hitch-hiking SFS is its long flat tail, consisting of mutations that fixed in the wavefront during the the sweep and were carried through much of the population. Intuitively, the underlying difference in the dynamics is that in a spatially structured population, an allele that begins to hitchhike midway through the sweep can reach very high frequencies, whereas in a well-mixed population hitchhiking is largely confined to those alleles present in the sweeping background very early in the sweep (Coop and Ralph 2012). Sweeps are also much slower in spatially structured populations than they are in well-mixed ones, giving mutation and recombination more time to act. We found that this makes the overall diversity higher, even at loci completely linked to the swept locus, and makes the intensity of hitchhiking decay more rapidly as one moves away from the swept locus along the genome. The high-frequency (flat) portion of the SFS relaxes back to the neutral expectation particularly quickly as genetic map distance to the swept locus increases.

The most pressing question is to what extent these effects might be seen in data from natural populations. For the restriction of hitchhiking to a narrow region of the genome, our results actually predict an absence of signal relative to a well-mixed population, so this would necessarily be present in natural populations with one-dimensional spatial structure. Indeed, Tavares *et al.* (2018) found that a pair of strong selective sweeps in a structured population of *Antirrhinum majus* reduced diversity only in narrow genomic windows. Since the width of the hitchhiking region on the genome is used to infer the selective coefficient of the sweep under the assumption that it is ≈*s*/ln(*Ns*), our results suggest that spatial structure should produce a bias towards underestimating the selective coefficients driving sweeps.

The question of whether the distinctive flat tail of the SFS should be visible in data is somewhat more subtle. Near any one swept locus, there will be very little diversity, so the full shape of the SFS will not be visible; the question is really whether there are likely to be *any* mutations found in this frequency range. Let us first consider the number of new mutations that are expected to hitchhike to high frequencies in essentially complete linkage with the sweeping allele. We predict that the expected SFS is *ξ*(*f*) ≈ *U*_n_ *T*_sweep_ in this regime. The regime extends over a range of frequencies of order 1, so *ξ* is also roughly the total expected number of alleles found in this frequency range. It might appear that the slower the sweep, the more alleles we should see, because of the factor of the sweep time *T*_sweep_ in *ξ*. But increasing *T*_sweep_ also decreases the length of the completely linked portion portion of the genome, which is ≈1/*T*_sweep_ in Morgans, reducing the mutation rate *U*_n_. Let *ν* be the mutation rate per Morgan, which is typically of order one. Then the locus-wide mutation rate will be *U*_n_ ≈ *ν*/*T*_sweep_. We see that the factors of *T*_sweep_ approximately cancel and the total expected number of high-frequency linked mutations is ξ ≈ *ν*, so it would be reasonable for many organisms to find such a mutation. Note that this is higher than the equivalent number for well-mixed populations by a factor ≈ ln(*Ns*), which might be an order of magnitude.

While one might expect to find a mutation or two in the high-frequency tail of the SFS around a sweep, one expects to find many more alleles that were introduced by recombination. At the characteristic map length *r* ≈ 1/*T*_sweep_ of the hitchhiking region, the recombinant alleles outnumber mutations that occurred during the sweep by a factor *U*_n_,eff/*U*_n_ ≈ *N*/*T*_sweep_ ≫ 1. (Here, *N* is really the neutral coalescence time *T*_coal_, i.e., the effective population size *Ne*.) This is mostly because each recombination event brings in a new haplotype that typically differs from the original sweeping background at multiple sites; specifically, over the map distance ≈ 1/*T*_sweep_ over which linkage disequilibrium is maintained during the sweep, they typically differ at ≈ *Nν*/*T*_sweep_ sites. If instead of alleles we count high-frequency recombinant haplotypes, we find that the expected number is order one, independent of any of the population parameters. Intuitively, this is because the limits of the region of the genome affected by hitchhiking are set by the map distance *r* ≈ 1/*T*_sweep_ at which we expect to find recombinants that successfully hitchhiked for long distances. In other words, the hitchhiking typically extends out to the first high-frequency recombinant in each direction on the genome from the swept locus. Thus, there will usually be a few high-frequency haplotypes, but not many more, as at larger map distances *rT*_sweep_ ≫ 1 recombination prevents haplotypes from hitchhiking to high frequencies. Here the contrast with well-mixed populations is stark: in a well-mixed population, hitchhiking is still broken up by successful recombinants occurring midway through the sweep, but there are typically a large number of these recombinants, each of which barely hitchhikes, and so only the original haplotype of the sweep reaches high frequency (Barton 1998; *Garud et al.* 2015).

Schrider *et al.* (2015) noted this phenomenon of potential high-frequency recombinant haplotypes around the selected locus, calling them the “soft shoulders” of the sweep and suggesting that they could potentially mislead inference methods into mistaking “hard” sweeps descended from a single mutation for “soft” sweeps descended from multiple independent mutations. But they only considered well-mixed populations and found that the soft shoulders could be reliably distinguished from the hard center by considering large genomic windows. It is unclear if this possible in one-dimensional populations, as the soft shoulders are much more frequent and much closer to the swept locus, as described above. It is possible that spatial structure may often lead to misidentification of hard sweeps as soft. While it is known that misspecified demography can interfere with sweep inference and distinguishing hard from soft sweeps (see, e.g., Harris *et al.* (2018)), the effect of space we have found here is particularly insidious because it is strong even in populations that appear to be only very weakly spatially structured by standard measures such as *F*_ST_.

Recent studies have found evidence for widespread soft sweeps and few hard sweeps in multiple species (*Garud et al.* 2015; Schrider and Kern 2017). However, most of these might better be termed “firm” sweeps, in that they must have begun from only a few mutations to be detectable; very soft sweeps with significant contributions from dozens or hundreds of mutations would not leave a strong enough trace. This is a surprising result in that it suggests some fine-tuning in nature. In standard population genetics models, a broad range of parameter space produces low beneficial mutation supplies and adaptation driven by hard sweeps, while a similarly broad range of parameter space produces abundant standing variation and adaptation via modest shifts in allele frequencies. Only a relatively narrow range of parameters reliably produces a few successful beneficial mutations. This suggests that either there must be some mechanism that drives populations to the right region of parameter space to produce firm sweeps, or the apparent prevalence of firm sweeps may be due to some other factor. For the first option, it may be that in populations with high levels of standing variation under selection, interference automatically reduces the number of fit backgrounds in such a way that the beneficial mutation supply effectively becomes of order one. However, to our knowledge there is as yet no model demonstrating this. Our results here suggest that spatial structure may be an example of the second option: a population feature which generically leads hard sweeps to appear firm.

There are a number of ways that the present work needs to be extended before we can fully assess the potential importance of spatial structure to hitchhiking in natural populations. We have focused on just the expected SFS, but (as shown by the extensive averaging that we need to do across simulation runs and allele frequencies; see Appendix A) in any particular sweep there will be large amounts of variation around the expectation, particularly at high frequencies. In addition, while have touched on potential haplotype structure here in the Discussion, we have not explicitly modeled haplotypes and linkage among neutral loci. This is most likely necessary to find patterns that can distinguish hard sweeps from soft sweeps in spatially structured populations; this may be a difficult problem, given that some of the features we find are reminiscent of spatial soft sweeps (Ralph and Coop 2010).

We have only considered a simple one-dimensional steppingstone model of spatial structure, but most natural populations are likely to be two-dimensional and have some long-range dispersal. Both of these are likely to reduce the effects of spatial structure on hitchhiking, as they increase the number of individuals that are contributing to the spread of the sweep at any point in time (Ralph and Coop 2010; Barton *et al.* 2013; Hallatschek and Fisher 2014; Fusco *et al.* 2016; Paulose and Hallatschek 2020). However, sweeps are still far slower, and the distribution of reproductive value across individuals carrying the sweeping allele is still far more skewed, than in a well-mixed population, so we expect some of the qualitative features of our results to persist. In particular, as long as the dispersal distribution is not too broad, it is still true that a single mutant or recombinant arising midway through the sweep can be the ancestor of a significant fraction of the population, so we still expect the SFS to have a flatter tail than the well-mixed *ξ*(*f*) ∝ 1/*f*^2^.

These extensions to our analysis are likely to be quite challenging analytically. For two-dimensional populations with purely short-range dispersal, the natural starting point would be Fusco *et al.* (2016)’s expressions for the distribution of “bubbles” and “sectors”, although one needs to check if they hold beyond the low-density limit. Alternatively, one could also try simply simulating hitchhiking under realistic parameter ranges for natural populations and measuring the resulting genetic diversity. Unfortunately, we know little about what the ranges of the relevant parameter values are in natural populations. The problem is particularly acute for the parameters related to spatial structure like the density *ρ*, the dispersal rate *m*, and the pattern of dispersal across space, especially the frequency and distribution of long-range jumps, which can determine the sweep dynamics (Ralph and Coop 2010; Hallatschek and Fisher 2014). This challenge could be potentially be addressed by first using genomic regions far from putative sweeps to infer the population structure, and then using this information to simulate sweeps.

## Data availability

All codes for simulation, Jupyter notebook for making the figures, and simulation data used in the Jupyter notebook are available at https://github.com/weissmanlab/SFS_spatial_sweep.

## Acknowledgements

The authors thank Ben Good and Adrian Gushin for helpful discussions. The computations in this paper were run on the FASRC Cannon cluster supported by the FAS Division of Science Research Computing Group at Harvard University.

## Appendix A. Simulation methods

As described in the Model section above, we use two-part simulations, first simulating the sweep forward in time, and then simulating coalescence at the neutral locus backward in time conditional on the sweep. The forward simulations are a standard stepping-stone Wright-Fisher model. Here, we will briefly explain the form of the backward simulations. At the start of each backward simulation, we have *n* lineages, sampled uniformly in space. We follow these backward through time, keeping track of their spatial location and which background they are on at the selected locus. (For the values of *n* used in each figure, see Table A1.) At the time of sampling, all are necessarily on the background of the sweeping allele, since the sweep has already been completed. Let *p*(*x*, *t*) be the sweep trajectory obtained from the forward simulation; going backward in time, the motion and coalescence of the lineages depends on this trajectory.

Each backward generation consists of three steps: dispersal, recombination, and coalescence, in that order. (Although because all rates are small, we do not expect the precise order to be important.) In the dispersal step, if a lineage is in the sweeping background in deme *x* at time *t* + 1, then at time *t* it will be in deme *x*, *x* - 1, or *x* + 1 with probabilities:

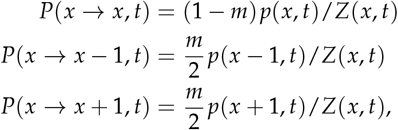

where *Z*(*x*, *t*) = *P*(*x → x*, *t*) + *P*(*x → x* - 1, *t*) + *P*(*x → x* + 1, *t*) is a normalization factor to ensure that the probabilities sum to one. For lineages on the ancestral background, the dispersal probabilities are the same but with *p* replaced by 1 - *p* in all the equations above. In the recombination step, each lineage that is in the sweeping background in deme *x* at time *t* (“after” dispersal) moves to the ancestral background with probability *r*(1 - *p*(*x*, *t*)), and each lineage in the ancestral background moves to the sweeping background with probability *rp*(*x*, *t*). In the coalescence step, two lineages that are both in deme *x* at time *t* coalesce with probability 1/(*ρp*(*x*, *t*)) if they are both on the sweeping background, or probability 1/(*ρ*(1 - *p*(*x*, *t*))) if they are both on the ancestral background. (If one is on the ancestral background and the other is on the sweeping background, they cannot coalesce.)

**Table A1.** Additional simulation parameter values. In all the listed figures, simulation results are averaged over a sliding window of width Δ *f* for all frequencies *f* > *f*_smooth_. We run *K*_forward_ independent forward simulations of the sweep. For each of these, we run *K*_backward_ independent backward simulations of the conditional coalescent process. All simulation results are the average over all these runs. Sample size *n* lists the number of sampled individuals at the start of each backward simulation.

| Figure | $f_{\text{smooth}}$ | $\Delta f$ | $K_{\text{forward}}$ | $K_{\text{backward}}$ | Sample size $n$ |
| --- | --- | --- | --- | --- | --- |
| 2 and 3, spatial | $10^{-3}$ | $3 \times 10^{-4}$ | $10^2$ | $10^3$ | $10^5$ |
| 2 and 3, well-mixed ($f < 8 \times 10^{-4}$) | $10^{-3}$ | $3 \times 10^{-4}$ | 10 | 2 | $10^5$ |
| 2 and 3, well-mixed ($f > 8 \times 10^{-4}$) | $10^{-3}$ | $3 \times 10^{-4}$ | $10^2$ | 10 | $10^4$ |
| 4 | $10^{-3}$ | $3 \times 10^{-4}$ | $10^2$ | $10^3$ | $10^5$ |
| 5 spatial | $7 \times 10^{-3}$ | $10^{-2}$ | $10^2$ | $10^3$ | $10^4$ |
| 5a well-mixed | $1.5 \times 10^{-2}$ | $2 \times 10^{-2}$ | $10^2$ | $10^2$ | $10^4$ |
| 5b-d well-mixed | $7 \times 10^{-3}$ | $10^{-2}$ | $10^2$ | $10^2$ | $10^4$ |
| C1 | $10^{-3}$ | $6 \times 10^{-4}$ | $5 \times 21$ | $10^3$ | $10^5$ |
| E1 | $10^{-3}$ | $3 \times 10^{-4}$ | $10^2$ | $10^3$ | $10^5$ |

For a locus completely linked to the selected locus, the entire sample must coalesce by the beginning of the sweep, since the sweep begins with a single mutant individual. For a locus that can recombine with the selected locus, some lineages may escape coalescence at the beginning of the sweep, in which case the sample will usually take far longer to fully coalesce, *T*_coal_ ~ *N*. Simulating this time in full detail would be prohibitive computationally, and wasteful, since over these long time scales the population is effectively well-mixed. Therefore, once the simulations reach times prior to the beginning of the sweep, *t* < 0, we continue them for 3200 generations to allow them to complete the “scattering phase” (Wakeley and Aliacar 2001), and then switch to a simple well-mixed Kingman coalescent until the sample is fully coalesced. For simulation parameters *m* = 0.25 and *L* = 500, these 3200 generations are less than the time *T*_mix_ ≈ 10^6^ required for lineages to disperse all the way across the range, but because the coalescence time scale *T*_coal_ ≈ *N* = 10^7^ is even longer we do not expect that this has a large impact on our results.

Rather than store full coalescent trees for our large samples, for each simulation we only keep track of the information we need to build the SFS across generations. Specifically, each generation we increment the total branch length ancestral to *k* sampled individuals, for all *k* from 1 to *n* - 1.

For every set of parameter values, we run *K*_forward_ independent forward simulations. (Here *K*forward is the number of successful simulations, not counting the many more where the beneficial mutant goes extinct without sweeping.) For each forward simulation, we run *K*_backward_ independent backward simulations. All simulation results presented are averages over these *K*_forward_ × *K*backward runs. The averaging over multiple backward simulations is particularly important, especially for estimating the spectrum at high frequencies *f*, and especially with recombination, as there can be substantial stochasticity in when the last few lineages coalesce. To deal with this further, in the figures we smooth the simulation curves by averaging over sliding windows of width Δ *f* for all frequencies *f* > *f*_smooth_. (Because the lower-frequency portions of the SFS receive many independent contributions from near the tips of each coalescent tree, they are already nearly deterministic and do not need additional smoothing.) The values of *K*_forward_, *K*_backward_, *f*_smooth_, and Δ *f* for each figure are specified in Table A1.

For the well-mixed SFS in Figure 2, we use two sets of simulations. For low frequencies *f* < 8 × 10^−4^ we need to use a large sample size *n*, but as mentioned above we need relatively few simulation runs. For higher frequencies, we can use a smaller sample size but must run more simulations. For Figure C1, we simulate 21 different possible starting locations for the sweep (ranging from *x* = 0 to *x* = 5000 in increments of 250). For each of these, we run five independent simulations. See Table A1 for the full simulation settings for both figures.

## Appendix B. Measuring wave speed

The speed of the wave of advance of the sweeping allele actually only reaches 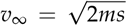 in infinitely dense populations. In real populations with finite density *ρ*, fluctuations reduce the speed of the wave (Brunet and Derrida 1997). For each value of *ρ*, we therefore measure *v* directly in a single forward simulation and use this value in all plots. To do this, we use Barton *et al.* (2013)’s definition of the wave speed: 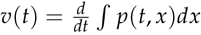. Since we are interested in the average speed, not the instantaneous speed of the wave front, we average *v*(*t*) over the middle half of the sweep, discarding the first *T*_sweep_/4 generations to avoid non-equilibrium effects while the wave is first establishing, and discarding the last *T*_sweep_/4 generations to avoid edge effects as the wave hits the far boundary of the range. We list the values we obtain for *v* in Table B1. They are all within ≈ 20% of the limiting value *v*_∞_, so these corrections are fairly minor.

## Appendix C. Sweeps starting from the middle of the range

If the genetic sweep starts somewhere in the middle of the range, the Fisher wave becomes bi-directional. Therefore, neutral alleles can fix in the left or the right wavefront. The sweep time also depends on the initial location of the beneficial mutation.

**Table B1.** Empirically measured wave speeds *v* that are used to plot the asymptotic behavior of the tail of SFS (*ξ*(*f*) = *U*_n_ *L*/*v* for Figures 2 and 4, *ξ*(*f*) = *U*_n_,eff *L*/*v* for Figure 5, and *ξ*(*f*) = 2*U*_n_ *L*(1 - *f*)/*v* for Figure C1). For comparison, the infinite-density limiting speed is 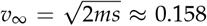 for the parameter values *m* = 0.25, *s* = 0.05 used in the simulations.

| Figure | $\rho$ | $v$ |
| --- | --- | --- |
| 2, 4 | $2 \times 10^4$ | 0.151 |
| 5 | $5 \times 10^3$ | 0.145 |
| C1 | 200 | 0.125 |

Suppose the beneficial mutation starts at spatial position *x* at *t* = 0. Then if a neutral mutation fixes in a wavefront at *t* = *t*_seed_, the allele frequency after the fixation is roughly *f* = (*x* - *vt*_seed_)/*L* if it is in the left side and *f* = (*L - l - vt*_seed_)/*L* if it on the right. There are different upper bounds for *t*_seed_ (0 ≤ *t*_seed_*x*/*v* for the left, 0 ≤ *t*_seed_(*L - x*)/*v* for the right) because the mutation has to fix before the wave arrives at the boundary. If the probability distribution of the starting location of the sweep is *q*(*x*), the expected high-frequency tail of the SFS is

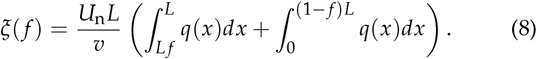

In the main text, we assume that the sweep starts in the leftmost deme (*q*(*x*) = *δ*(*x*)), and therefore get *ξ*(*f*) = *U*_n_ *L*/*v*. If the sweep starts at some other position *l*, which we can assume without loss of generality to be < *L*/2, then *q*(*x*) = *δ*(*x* - *l*) and we have:

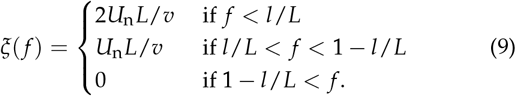

If we instead consider the SFS averaged over the genomic neighborhoods of many independent sweeps with starting positions uniformly distributed over the range, we have *q*(*x*) = 1/*L* and *ξ*(*f*) = (2*U*_n_ *L*/*v*)(1 - *f*), as shown in Figure C1.

## Appendix D. Deviation from the uniform distribution at very high frequencies

Neutral mutations are unlikely to reach very high frequencies *f* → 1 because they take a finite amount of time to fix in the wavefront, and during this time the wave advances. We can therefore estimate at which frequency the simulated expected SFS starts deviating from the uniform distribution, *ξ*(*f*) = *U*_n_ *L*/*v* by considering how long it takes for a lineage to take over the wavefront. Using Barton *et al.* (2013)’s approximation for the wavefront coalescence time, 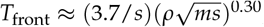 a neutral mutation is very unlikely to fix in the first fraction *vT*front/*L* of the range. In other words, the uniform tail of the SFS should be cut off at *f* ≈ 1 - *vT*front/*L*. For the parameter values in Figure 2, this is *f* ≲ 1 - *vT*_front_/*L* ≈ 0.8, which roughly agrees with the simulation results.

## Appendix E. Site frequency spectrum from surfing mutations

We want to find an approximate expression for the SFS at intermediate frequencies where it is primarily composed of alleles that surf in the wavefront before being dropped (Figure 3). To do this, we follow the wavefront in the co-moving frame, viewing it as a small population of size 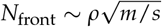. If we imagine tracing out the trajectory of the frequency of the allele in the wavefront over time, the total frequency in the entire population will be proportional to the area under this curve (Okada and Hallatschek 2021). Specifically, suppose that the allele persists in the wavefront for *t* generations at a frequency of ≈ *y* before being dropped. The total number of copies of the allele will then be ~ *vtyρ*, for a total frequency of *f* ~ *vtyρ*/(*ρL*) = *vty*/*L*.

**Figure C1.**
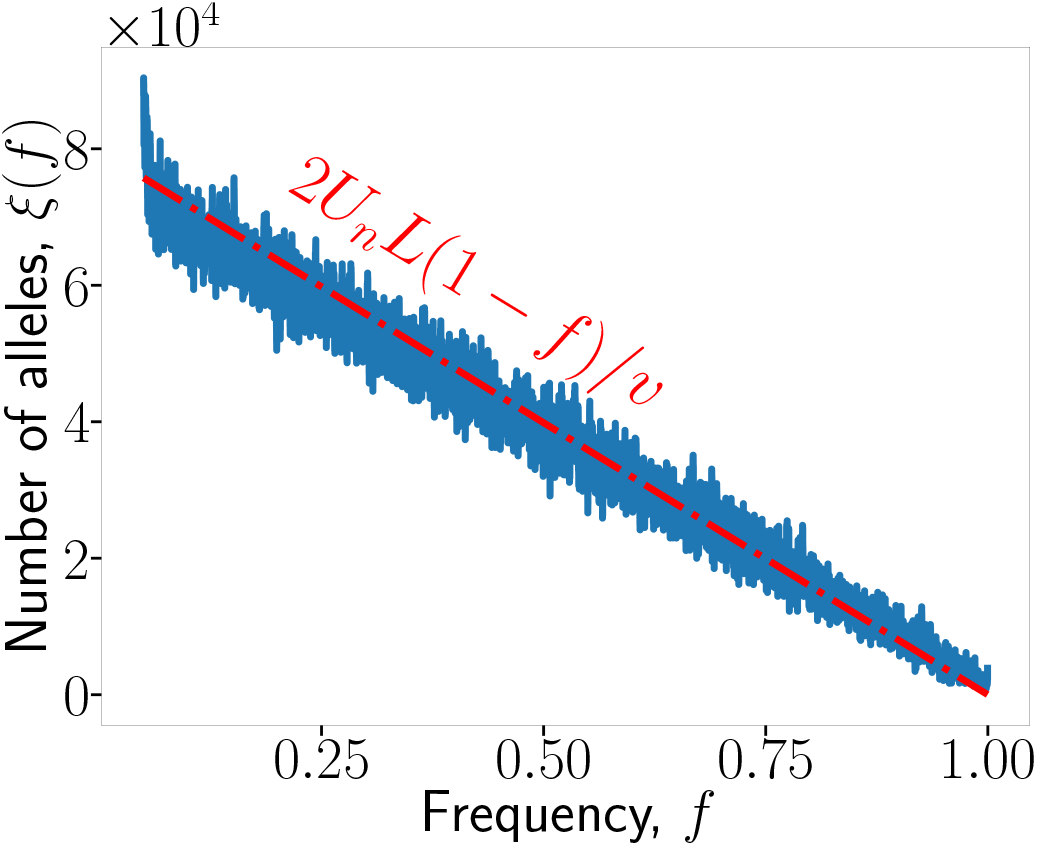
The tail of an expected SFS when the origin of the sweep is uniformly sampled from *x* ∊ [0, *L*). *L* = 5001, *ρ* = 2 × 10^2^, *m* = 0.25, *s* = 0.05, *U*_n_ = 1. SFS is measured immediately after the end of the sweep.

We now need to approximate the joint distribution of the frequency *y* that an allele reaches in the wavefront and the time *t*(*y*) for which it surfs conditional on reaching that frequency. Coarse-graining over the time scale of coalescence in the wave-front (roughly, the time ~ 1/*s* for it to travel its own width, up to logarithmic factors (Barton *et al.* 2013; Birzu *et al.* 2018)), we can treat the wavefront as having an effective “sweepstakes” offspring distribution with tail exponent 1 (Okada and Hallatschek 2021). The typical persistence time then is short and only logarithmically dependent on *y* (Okada and Hallatschek 2021). The exact form of this logarithmic dependence is unclear though, and most likely it only describes extremely dense populations (Birzu *et al.* 2018; Barton *et al.* 2013). Instead, we will follow Barton *et al.* (2013) and Birzu *et al.* (2018) in approximating it with a power law. Okada and Hallatschek (2021) find that for sweepstakes reproduction with tail exponent > −1, the overall coalescence time of the population scales as a power of the population size, *T* ∝ *N^β^* (Schweinsberg 2003), and the typical time for a lineage to reach *n* < *N* copies follows the same power law, *t* ∝ *n^β^*. We guess that we can extend this pattern to tail exponent = −1. Then Barton *et al.* (2013)’s expression 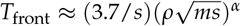 for the overall coalescence time means that the typical persistence time for a lineage that reaches wave-front frequency *y* should be 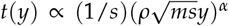, with the same exponent *α*. With this guess, we can rewrite the allele’s overall frequency *f* as a function of its wavefront frequency *y*:

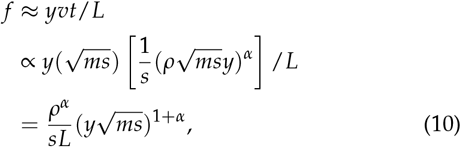

up to *O*(1) numerical factors.

To find the probability *P*(*f*) that an allele reaches overall frequency *f*, note that the probability that the allele reaches a wavefront frequency of at least *y* is just the standard *P*(*y*) ≈ 1/(*yN*_front_) (Okada and Hallatschek 2021). Inverting Eq. (10) to find *y* in terms of *f* then gives 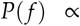 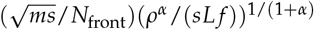. Differentiating with respect to *f*, we find the probability density *p*(*f*):

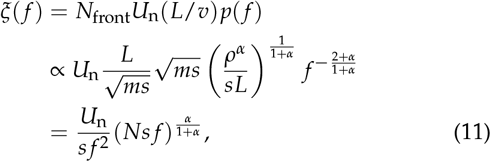

again ignoring *O*(1) numerical factors. From the density, we can immediately obtain the site frequency spectrum *ξ*(*f*) by multiplying by the total number of mutations that occur in the wavefront, which is the product of the wavefront mutation supply *N*_front_*U*_n_ and the sweep time *T*_sweep_ = *L*/*v*:

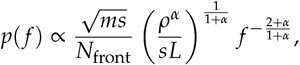

where in the last line we have used that the total population size is *N* = *Lρ*. We have neglected numerical constants throughout this argument, so there is an undetermined constant of proportionality. Comparing Eq. (11) to simulations, it appears that this constant is ≈ 0.6 (Figure 3).

To check the accuracy of our guesses and approximations, we test how well Eq. (11) fits simulations over a range of parameter values. While computational limitations prevent us from varying the parameters over even an order of magnitude, the prediction does appear to match well (Figure E1).

**Figure E1.**
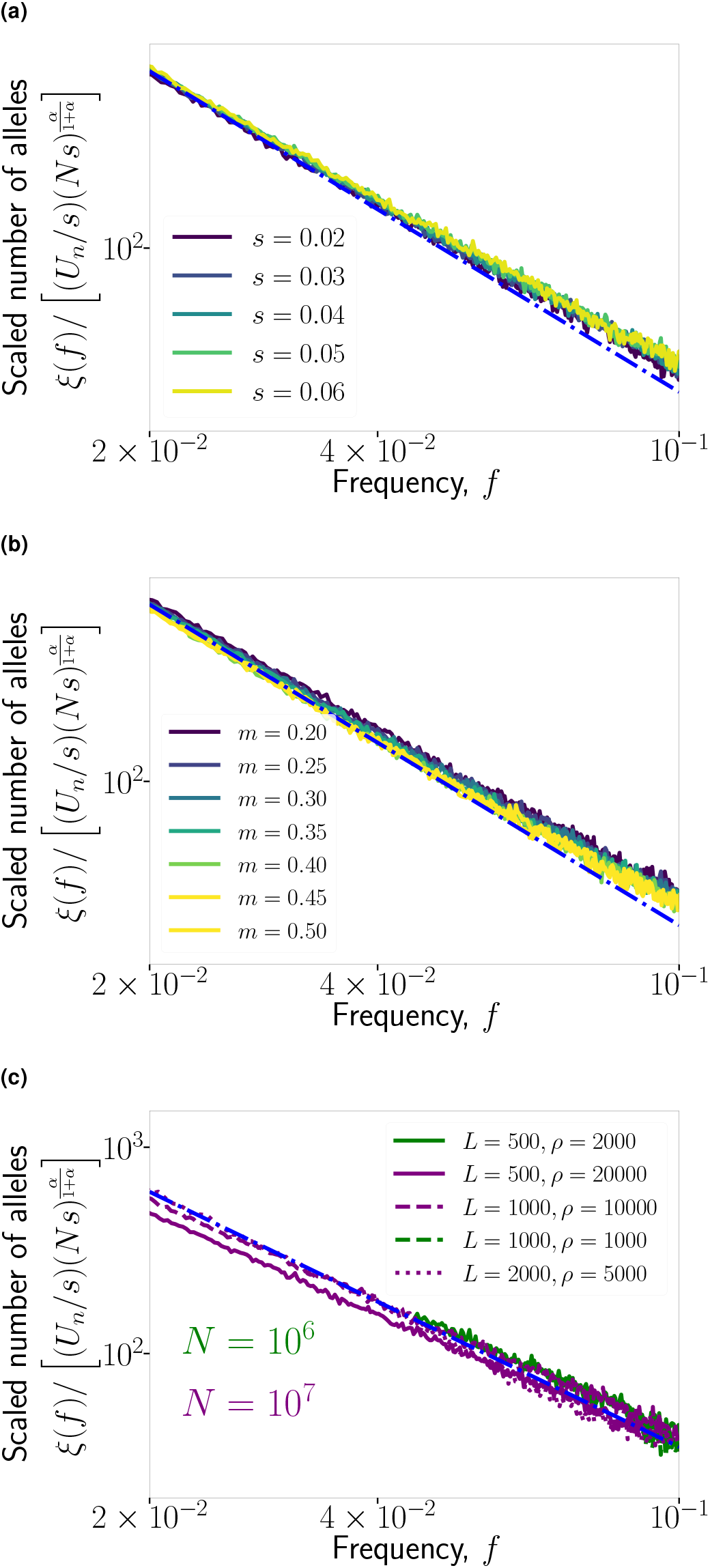
The analytical prediction Eq. (11) for the SFS produced by surfing alleles matches simulations over a range of parameter values. All plots show spectra rescaled by the factor that Eq. (11) predicts will collapse the results. The blue dot-dashed line is the rescaled version of the prediction shown in Figure 3, i.e., 0.6/ *f*^(2+*α*)/(1+*α*)^. All other curves are simulation results. If not specified otherwise, the parameter values are set to *L* = 5 × 10^2^, *ρ* = 2 × 10^4^, *s* = 0.05, *m* = 0.25, *r* = 0. All spectra are measured immediately after the completion of the sweep.

## Literature Cited

Allman, B. E. and D. B. Weissman, 2018 Hitchhiking in space: Ancestry in adapting, spatially extended populations. Evolution 72: 722–734.

Barton, N. H., 1998 The effect of hitch-hiking on neutral genealogies. Genetical Research 72: 123–133.

Barton, N. H., 2000 Genetic hitchhiking. Philosophical Transactions of the Royal Society B: Biological Sciences 355: 1553–1562.

Barton, N. H., A. M. Etheridge, J. Kelleher, and A. Véber, 2013 Genetic hitchhiking in spatially extended populations. Theoretical population biology 87: 75–89.

Birzu, G., O. Hallatschek, and K. S. Korolev, 2018 Fluctuations uncover a distinct class of traveling waves. Proceedings of the National Academy of Sciences 115: E3645–E3654.

Bisschop, G., K. Lohse, and D. Setter, 2021 Sweeps in time: leveraging the joint distribution of branch lengths. bioRxiv p. 2021.01.27.428367.

Booker, T. R., S. Yeaman, and M. C. Whitlock, 2020 Global adaptation complicates the interpretation of genome scans for local adaptation. Evolution Letters.

Bourgeois, Y. X. C. and B. H. Warren, 2021 An overview of current population genomics methods for the analysis of whole-genome resequencing data in eukaryotes. Molecular Ecology 30: 6036–6071.

Brunet, E. and B. Derrida, 1997 Shift in the velocity of a front due to a cutoff. Physical Review E 56: 2597–2604.

Coop, G., J. K. Pickrell, J. Novembre, S. Kudaravalli, J. Li, et al., 2009 The role of geography in human adaptation. PLOS Genetics 5: e1000500.

Coop, G. and P. Ralph, 2012 Patterns of neutral diversity under general models of selective sweeps. Genetics 192: 205–224.

DeGiorgio, M., C. D. Huber, M. J. Hubisz, I. Hellmann, and R. Nielsen, 2016 Sweepfinder2: increased sensitivity, robustness and flexibility. BIOINFORMATICS 32: 1895–1897.

Desai, M. M. and D. S. Fisher, 2007 Beneficial mutation selection balance and the effect of linkage on positive selection. Genetics 176: 1759–1798.

Fay, J. C. and C.-I. Wu, 2000 Hitchhiking under positive Darwinian selection. Genetics 155: 1405–1413.

Fisher, R. A., 1937 The wave of advance of advantageous genes. Annals of Eugenics 7: 355–369.

Fusco, D., M. Gralka, J. Kayser, A. Anderson, and O. Hallatschek, 2016 Excess of mutational jackpot events in expanding populations revealed by spatial Luria–Delbrück experiments. Nature communications 7: 1–9.

Garud, N. R., P. W. Messer, E. O. Buzbas, and D. A. Petrov, 2015 Recent selective sweeps in North American *Drosophila melanogaster* show signatures of soft sweeps. PLoS genetics 11: e1005004–e1005004.

Gillespie, J. H., 2000a Genetic drift in an infinite population: the pseudohitchhiking model. Genetics 155: 909–919.

Gillespie, J. H., 2000b The neutral theory in an infinite population. Gene 261: 11–18.

Hallatschek, O. and D. S. Fisher, 2014 Acceleration of evolutionary spread by long-range dispersal. Proceedings of the National Academy of Sciences 111: E4911–E4919.

Harris, A. M., N. R. Garud, and M. DeGiorgio, 2018 Detection and classification of hard and soft sweeps from unphased genotypes by multilocus genotype identity. Genetics 210: 1429–1452.

Hartl, D. L., 2020 A primer of population genetics and genomics. Oxford University Press.

Hartl, D. L. and A. G. Clark, 1997 Principles of population genetics, volume 116. Sinauer.

Hejase, H. A., N. Dukler, and A. Siepel, 2020 From summary statistics to gene trees: Methods for inferring positive selection. Trends in Genetics 36: 243–258.

Hermisson, J. and P. S. Pennings, 2005 Soft sweeps: Molecular population genetics of adaptation from standing genetic variation. Genetics 169: 2335–2352.

Kaplan, N. L., R. R. Hudson, and C. H. Langley, 1989 The “hitchhiking effect” revisited. Genetics 123: 887–899.

Karasov, T., P. W. Messer, and D. A. Petrov, 2010 Evidence that adaptation in *Drosophila* is not limited by mutation at single sites. PLoS Genetics 6: e1000924.

Kern, A. D. and D. R. Schrider, 2018 diploS/HIC: An updated approach to classifying selective sweeps. G3 8: 1959–1970.

Kim, Y. and T. Maruki, 2011 Hitchhiking effect of a beneficial mutation spreading in a subdivided population. Genetics 189: 213–226.

Kim, Y. and W. Stephan, 2002 Detecting a local signature of genetic hitchhiking along a recombining chromosome. Genetics 160: 765–777.

Li, H. and W. Stephan, 2006 Inferring the demographic history and rate of adaptive substitution in *Drosophila*. PLoS Genetics 2: e166.

Luria, S. E. and M. Delbrück, 1943 Mutations of bacteria from virus sensitivity to virus resistance. Genetics 28: 491.

Maruyama, T., 1971 An invariant property of a structured population. Genet Res 18: 81–84.

Maynard Smith, J. and J. Haigh, 1974 The hitch-hiking effect of a favourable gene. Genetical Research 23: 23–35.

Okada, T. and O. Hallatschek, 2021 Dynamic sampling bias and overdispersion induced by skewed offspring distributions. Genetics 219: iyab135.

Paulose, J. and O. Hallatschek, 2020 The impact of long-range dispersal on gene surfing. Proceedings of the National Academy of Sciences - PNAS 117: 7584–7593.

Pavlidis, P., D. Zivkovic, A. Stamatakis, and N. Alachiotis, 2013 Sweed: Likelihood-based detection of selective sweeps in thousands of genomes. Molecular biology and evolution 30: 2224–2234.

Ralph, P. L. and G. Coop, 2010 Parallel adaptation: One or many waves of advance of an advantageous allele? Genetics 186: 647–668.

Sabeti, P. C., P. Varilly, B. Fry, J. Lohmueller, E. Hostetter, et al., 2007 Genome-wide detection and characterization of positive selection in human populations. Nature 449: 913–918.

Sattath, S., E. Elyashiv, O. Kolodny, Y. Rinott, and G. Sella, 2011 Pervasive adaptive protein evolution apparent in diversity patterns around amino acid substitutions in *Drosophila simulans*. PLoS Genetics 7: e1001302.

Schrider, D. R. and A. D. Kern, 2016 S/hic: Robust identification of soft and hard sweeps using machine learning. PLoS genetics 12: e1005928–e1005928.

Schrider, D. R. and A. D. Kern, 2017 Soft sweeps are the dominant mode of adaptation in the human genome. Molecular biology and evolution 34: 1863–1877.

Schrider, D. R., F. K. Mendes, M. W. Hahn, and A. D. Kern, 2015 Soft shoulders ahead: spurious signatures of soft and partial selective sweeps result from linked hard sweeps. Genetics 200: 267–284.

Schweinsberg, J. R., 2003 Coalescent processes obtained from supercritical Galton-Watson processes. Stochastic Processes and their Applications 106: 107–139.

Slatkin, M. and T. Wiehe, 1998 Genetic hitch-hiking in a subdivided population. Genetics Research 71: 155–160.

Smith, J., G. Coop, M. Stephens, and J. Novembre, 2018 Estimating time to the common ancestor for a beneficial allele. Molecular Biology and Evolution 35: 1003–1017.

Stephan, W., 2019 Selective sweeps. Genetics 211: 5–13.

Stephan, W., T. H. E. Wiehe, and M. W. Lenz, 1992 The effect of strongly selected substitutions on neutral polymorphism: analytical results based on diffusion theory. Theoretical Population Biology 41.

Stern, A. J., P. R. Wilton, and R. Nielsen, 2019 An approximate full-likelihood method for inferring selection and allele frequency trajectories from DNA sequence data. PLOS Genetics 15: e1008384.

Tang, K., K. R. Thornton, and M. Stoneking, 2007 A new approach for using genome scans to detect recent positive selection in the human genome. PLoS biology 5: e171.

Tavares, H., A. Whibley, D. L. Field, D. Bradley, M. Couchman, et al., 2018 Selection and gene flow shape genomic islands that control floral guides. Proceedings of the National Academy of Sciences 115: 11006–11011.

Vitti, J. J., S. R. Grossman, and P. C. Sabeti, 2013 Detecting natural selection in genomic data. Annual review of genetics 47: 97–120.

Wakeley, J., 2008 Coalescent Theory: An Introduction. Macmillan.

Wakeley, J. and N. Aliacar, 2001 Gene genealogies in a metapopulation. Genetics 159: 893–905.

Weissman, D. B. and N. H. Barton, 2012 Limits to the rate of adaptive substitution in sexual populations. PLoS genetics 8: e1002740.

